# BERBERINE BRIDGE ENZYME-LIKE OXIDASES OF CELLODEXTRINS AND MIXED-LINKED β-GLUCANS CONTROL SEED COAT FORMATION

**DOI:** 10.1101/2023.02.24.529966

**Authors:** Sara Costantini, Manuel Benedetti, Daniela Pontiggia, Moira Giovannoni, Felice Cervone, Benedetta Mattei, Giulia De Lorenzo

**Affiliations:** Dipartimento di Biologia e Biotecnologie “C. Darwin”, Sapienza University of Rome; Department of Life, Health and Environmental Sciences, University of L’Aquila, 67100 L’Aquila, Italy

**Author notes:** These authors equally contributed to this work (Sara Costantini, Manuel Benedetti).

**Keywords:** berberine bridge enzyme-like oxidases, cellodextrins, mixed-linked β-glucans, damage-associated molecular patterns, seed coat

## Abstract

A member of the Arabidopsis Berberine Bridge Enzyme-like (BBE-l) protein family named CELLODEXTRIN OXIDASE 2 (CELLOX2) has been characterized in this paper and shown to display structural and enzymatic features similar to the previously characterized CELLOX1. These include the capability to oxidize the mixed-linked β-1→3/β-1→4-glucans (MLGs), recently described as cell wall-derived damage-associated molecular patterns (DAMPs) that activate plant immunity. The two paralogous genes show a different expression profile. Unlike *CELLOX1, CELLOX2* is not expressed in seedlings or in adult plants and is not involved in immunity against *Botrytis cinerea.* Both genes are expressed in a concerted manner in the seed coat during development: whereas *CELLOX2* transcripts are detected mainly during the heart stage, *CELLOX1* transcripts are detected later, when the expression of *CELLOX2* decreases. Analysis of seeds of *cellox1* and *cellox2* knock-out mutants show alterations in the structure of the coat and mucilage, but not in their monosaccharide composition. We propose that the cell wall structure of specific organs is not only the result of a coordinated synthesis/degradation of polysaccharides but also of their exposure to enzymatic oxidation. Our results also reinforce the view that the family of BBE-l proteins is at least in part devoted to the control of the activity of cell wall-derived oligosaccharides acting as DAMPs.

**SENTENCE:** Two Arabidopsis BBE–like oxidases of the cell wall DAMPs cellodextrins and mixed-linked β-glucans inactivate their elicitor activity. Seed coat and mucilage are altered in null mutants of two enzymes.

## INTRODUCTION

Sensing the presence of microbe-associated molecular patterns (MAMPs) and damage-associated molecular patterns (DAMPs) alerts the plants on the presence of dangerous situations and is crucial for immunity and survival (De Lorenzo and Cervone, 2022). DAMPs include several fragments of the plant cell wall such as oligogalacturonides (OGs) and cellodextrins (CDs) (Pontiggia et al., 2020) as well as fragments of xyloglucan (Claverie et al., 2018), mannan (Zang et al., 2019), arabinoxylan (Mélida et al., 2020) and mixed-linked β-1→3/β-1→4-glucan (MLGs) (Barghahn et al., 2021; Rebaque et al., 2021; Yang et al., 2021). These molecules induce typical responses of the pattern-triggered immunity (PTI) generally induced by MAMPs. It is reported that DAMPs such as OGs and CDs are under homeostatic control in Arabidopsis, being deactivated, respectively, by four specific OG oxidases named OGOX1-4 (Benedetti et al., 2018) and one CD oxidase named CELLOX (Locci et al., 2019). OGOXs and CELLOX belong to the large family of FAD-binding berberine-bridge enzyme-like (BBE-l) proteins (Daniel et al., 2017). The physiological role of many oligosaccharide oxidases belonging to BBE-l family is still unexplored (Pontiggia et al., 2020). The substrates of the Arabidopsis CELLOX, encoded by the gene *BBE22/At4g20860*, are CDs with a DP higher than 2, markedly cellotriose, which is the most efficient elicitor-active fragment of cellulose (Locci et al., 2019). Remarkably, monomeric glucose is not a substrate of CELLOX, indicating that the specificity of this enzyme is different from that of two other carbohydrate oxidases, i.e. the *Helianthus annuus* carbohydrate oxidase Ha-CHOX and the *Lactuca sativa* carbohydrate oxidase Ls-CHOX, also belonging to the BBE-l family (Custers et al., 2004). Oxidation of CDs by CELLOX occurs at the C1 of the residue at the reducing end and produces gluconic acid and H_2_O_2_, i.e., a molecule locally formed as a consequence of cellulose damage that can potentially act as a positional signal in cell-to-cell communication (Pontiggia et al., 2020).

Both OGOXs and CELLOX are apoplastic proteins and display a markedly alkaline pH optimum (Benedetti et al., 2018; Locci et al., 2019), being the alkalinization of the apoplast one of the early plant responses to microbial threats (Felle et al., 2005). The over-expression of OGOX1 and CELLOX in transgenic Arabidopsis plants leads to an increased resistance against infection by *Botrytis cinerea*, likely because these enzymes impede an efficient digestion of the oxidated pectin and cellulose fragments by the fungal degrading enzymes (Benedetti et al., 2018; Locci et al., 2019).

*CELLOX* is present on chromosome 4, in the same cluster that includes *OGOX1* and *OGOX2* and shows a remarkably similar expression profile with *OGOX1* during the immune response (Locci et al., 2019). The closest paralogue of *CELLOX/BBE22/At4g20860*, i.e. *At5g44360/BBE23*, is characterized in this work and shown to encode a protein active on CDs and with a similar specificity to the previously characterized CELLOX, which from now on is renamed CELLOX1 whereas BBE23 is named CELLOX2. We report here that both enzymes display oxidative activity not only on CDs but also on MLGs and have a different pattern of expression and distinct roles since, unlike CELLOX1, CELLOX2 is not involved in immunity of Arabidopsis against *B. cinerea*. Both CELLOX1 and CELLOX2 instead influence the coat structure and mucilage porosity of seeds during their development. This paper introduces the notion that the cell wall structure in some organs and tissues may be influenced not only by the synthesis and degradation of the polysaccharides but also by their exposure to enzymatic oxidation. This paper also reinforces the notion that some DAMPs may be involved both in immunity and development.

## RESULTS

### BBE22 and BBE23 are oxidases of both cellodextrins and MLGs

The closest paralogue of the gene *CELLOX/At4g20860/BBE22* in Arabidopsis is the 1599 bp gene *At5g44360* also described as *BBE23* (Data S1), the product of which is still uncharacterized in terms of activity and physiological role. *At5g44360/BBE23* is localized on chromosome 5, in a gene cluster that includes five other paralogues (*At5g44380/BBE24, At5g44390/BBE25, At5g44400/BBE26, At5g44410/BBE27, At5g44440/BBE28).* The *At5g44360/BBE23*-encoded product (BBE23) is a 532 amino acid-long protein with an expected molecular mass of 60.49 kDa, and shares with BBE22 65.21% of amino acid identity and the typical oxygen binding motif PTVGVGG (Locci et al., 2019; Scortica et al., 2022). A homology-based molecular modelling of the 3D structure using a template the crystallographic structure of the monolignol oxidase BBE15 encoded by *At2g34790* (code 4UD8 in the Protein Data Bank, http://www.pdb.org) shows that BBE23 shares with BBE22 the cysteine and histidine residues to bind FAD and two tyrosine residues required for the proton abstraction, typical features of the type IV active site (Figure S1) (Daniel et al., 2017; Messenlehner et al., 2021). The same residues are also present in the PpCBOX of cellobiose oxidase from *Physcomitrella patens* (Toplak et al., 2018). At the physiological acidic pH of the apoplast, BBE23 shares with BBE22 the negatively charged residues surrounding the active site (Locci et al., 2019). Both BBE22 and BBE23 maintain the typical oxygen gate-keeper valine (V157) and the surrounding residues (Locci et al., 2019) and, according to SignalP-5.0 (https://services.healthtech.dtu.dk/service.php?SignalP-5.0) and Uniprot (www.uniprot.org), are predicted to be extracellular proteins. Coherently with the prediction, the fluorescence of the transiently expressed GFP-tagged form of BBE23, agroinfiltrated in *Nicotiana tabacum* leaves and analyzed by confocal microscopy, was localized mainly in the cell periphery that includes the apoplast (Figure 1).

**Figure 1.**
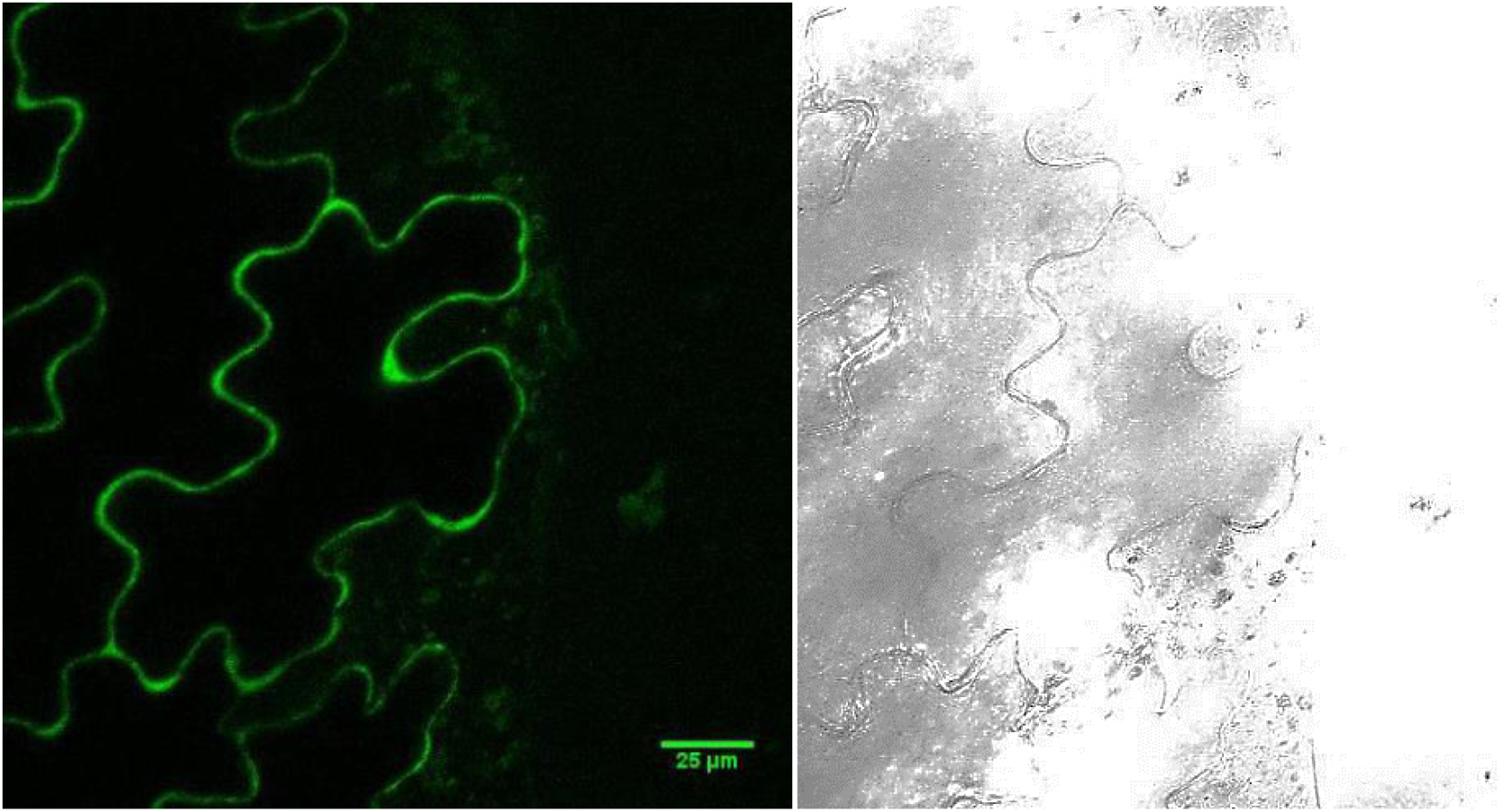
Transient expression of 35S::*CELLOX2-GFP* in agro-infiltrated tobacco leaves. The distribution of CELLOX2-GFP fluorescence at the cell periphery is in agreement with a localization in the apoplast. Confocal microscopy image (left); bright field (right).

BBE23 fused downstream a 6xHis-tag (H-BBE23) and a Flag-His-SUMO-Star tag (FHS-BBE23) were expressed as secreted proteins in *Pichia pastoris* for biochemical characterization (Figure 2A and Data S2). Immunoblot analysis of methanol-induced culture filtrates of *P. pastoris* revealed that FHS-BBE23 is expressed at low levels, whereas H-BBE23 is undetectable (Figure 2B). The culture filtrate from *P. pastoris* expressing FHS-BBE23 was analyzed in parallel to the culture filtrates of *P. pastoris* expressing FHS-BBE22 (Scortica et al., 2021; Scortica et al., 2022). Both proteins are expressed as highly glycosylated forms since they appeared as a smeared signal and, after treatment with PNGase F, as a single band of 75 kDa (Figure 2C). FHS-BBE22 and FHS-BBE23 were purified as reported (Scortica et al., 2022) with a protein yield ranging from 0.1 mg.L^-1^ (FHS-BBE23) to 0.5 mg.L^-1^ (FHS-BBE22) (Figure 3A). The activity of pure FHS-BBE22 and FHS-BBE23 was compared by measuring the amount of H_2_O_2_ produced in the presence of different substrates (Figure 3B). The proteins were not active towards OGs, chito-oligosaccharides, maltose and D-lactose as well as the monosaccharides D-glucose, D-xylose, D-galactose and N-acetyl-glucosamine. Instead, activity was observed using CDs of different length as substrates and was higher for oligomers with a degree of polymerization ≥ 3 (Figure 3B). BBE23 was, therefore, named CELLODEXTRIN OXIDASE 2 (CELLOX2), whereas BBE22, previously named CELLOX (Locci et al., 2019) was renamed as CELLOX1.

**Figure 2.**
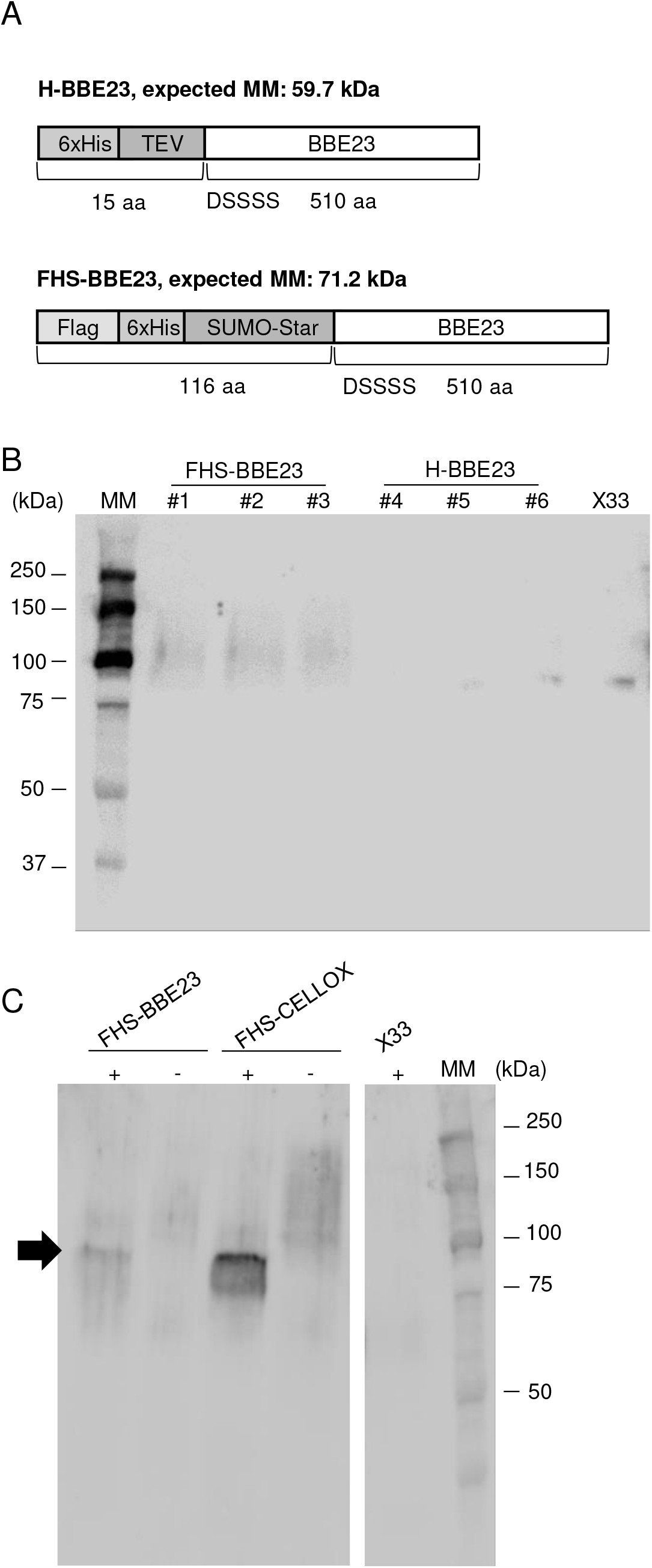
Heterologous expression of BBE23 in *P. pastoris.* A) Schematic representation of His-tagged (top) and Flag-His-SUMO-Star-tagged (bottom) version of BBE23, referred to as H-BBE23 and FHS-BBE23, respectively. The amino acid sequence of mature BBE23 (starting with DSSSS) was fused downstream of the His-TEV- and Flag-His-SUMO-Star-tag. B) Immuno-decoration analysis on the methanol-induced culture filtrates from different transformants expressing FHS-BBE23 (#1-3) and H-BBE23 (#4-6). C) Immuno-decoration analysis on untreated (-) and PNGase F-treated (+, deglycosylated) culture filtrates from the two highest expressing FHS-CELLOX and FHS-BBE23 transformants. The black arrow points to the PNGase F-treated FHS-BBE23. Molecular weight marker (MM) is also shown in (B) and (C).

**Figure 3.**
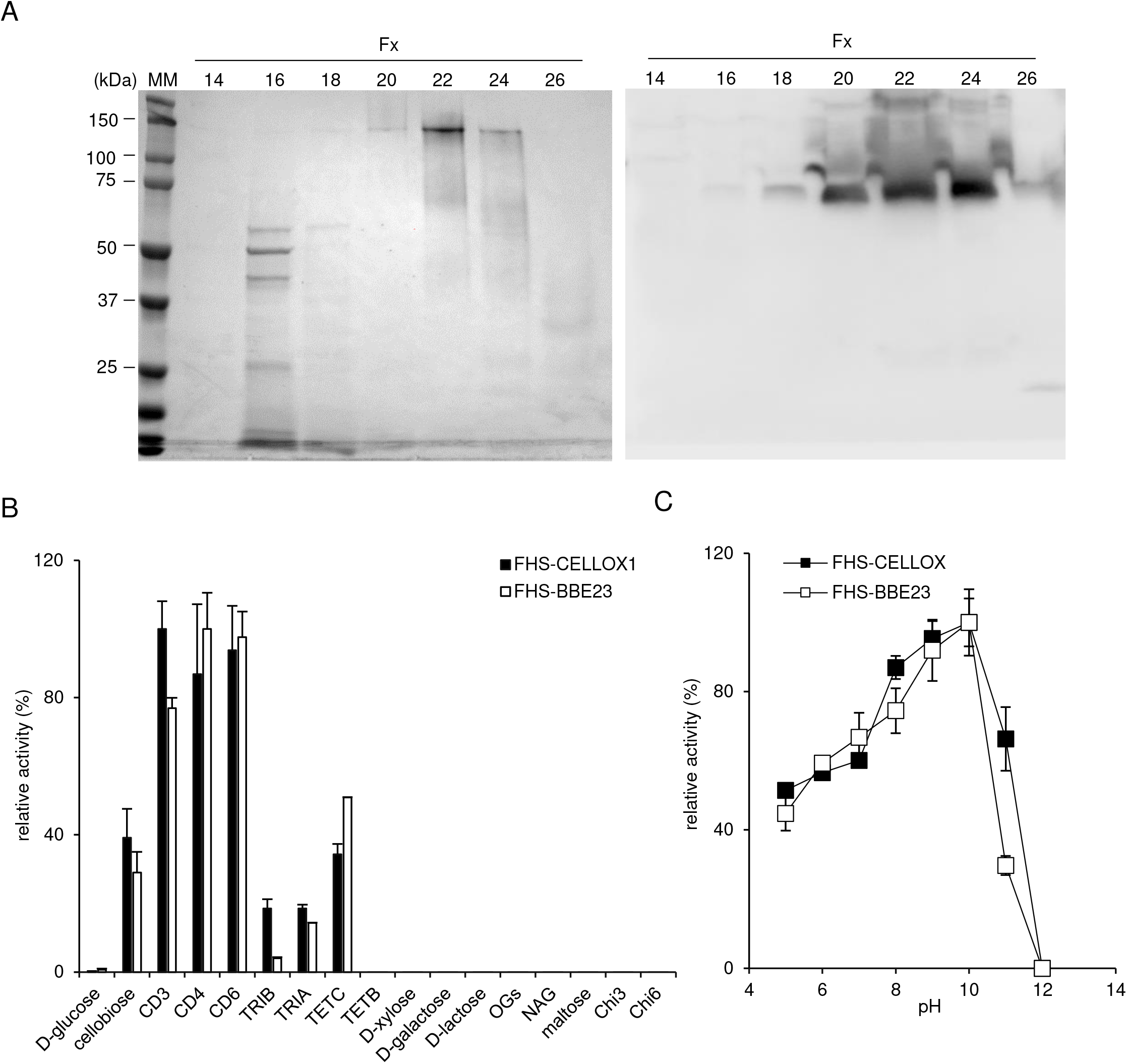
Biochemical characterization of FHS-BBE23. A) *Left*, SDS-PAGE/Coomassie blue staining analysis of fractions (fx) eluted from the 6xHistidine tag-affinity chromatography and used for the biochemical characterization (fx 20-25). *Right*, Immuno-decoration analysis of the same fractions shown in the left panel upon treatment with PNGase F. Molecular weight marker (MM) is also shown. B) Relative activity of FHS-BBE23 and FHS-CELLOX (flag-his-sumoylated form of the enzymes) towards different substrates as determined at pH 7.5 by the xylenol-orange assay. C) pH-dependent activity of FHS-BBE23 and FHS-CELLOX using CD2 as substrate. [CD2, cellobiose; CD3, cellotriose; CD4, cellotetraose; CD6, cellohexaose; Chi3, chitotriose; Chi6, chitohexaose; TETB, 3-β-D-cellotriosyl-glucose; TETC, mixture of 3-β-D-cellobiosyl-cellobiose and 3-β-D-glucosyl-cellotriose; TRIA, 3-β-D-glucosyl-cellobiose; TRIB, 3-β-D-cellobiosyl-glucose; OGs, oligogalacturonides; NAG, N-acetyl-glucosamine]

The optimal activity of CELLOX2, determined by using CD2 as substrate, was detected at pH 10 (Figure 3C), a markedly alkaline pH similar to the optimal pH of CELLOX1 and other BBE-like oligosaccharide oxidases such as OGOX1 (Benedetti et al., 2018; Scortica et al., 2021). It is likely that, at the alkaline pH, the electrostatic potential surface surrounding the active site of CELLOXs becomes more negative, possibly facilitating the interaction with the respective substrate.

Notably, both CELLOX1 and CELLOX2 showed activity on MLGs, which have been reported to act as DAMPs (Barghahn et al., 2021; Rebaque et al., 2021; Yang et al., 2021). Among the different MLGs tested, the highest activity was observed on a mixture of 3-β-D-cellobiosyl-cellobiose and 3-β-D-glucosyl-cellotriose (TETC), followed by 3,1-β-D-cellobiosyl-glucose (TRIB) and 3-β-D-glucosyl-cellobiose (TRIA), whereas no activity was detected in the presence of 3-β-D-cellotriosyl-glucose (TETB) (Figure 3B). The MLG-oxidizing activity of CELLOX1 was further analyzed and shown to be similar to the MLG-oxidizing activity of CELLOX2 (Figure S2). At pH 8.5, CELLOX1 was active on TRIB (0.12 μmol H_2_O_2_ min^-1^ mg^-1^), TRIA (0.14 μmol H_2_O_2_ min^-1^ mg^-1^) and TETC (0.28 μmol H_2_O_2_ min^-1^ mg^-1^). These values were about 2-4-fold lower than those measured on substrates such as CD3 and CD4 (Figure S2). However, the chromatographic profiles of the HPAEC-PAD analysis showed that the commercial TRIB contained relevant contaminating amounts of CD2 (40%) and CD3 (4%) (Figure S3), whereas TRIA and TETC were substantially pure (Figure 4A and 4C). Therefore, we concluded that the enzyme activity observed by using TRIB as substrate (Figure 3B, Figure S2) was essentially due to the oxidation of the contaminants CD2 and CD3 (Figure S3), and that CELLOX1 oxidizes specifically TRIA and TETC (Figure 4B e 4C). MS analyses showed that the oxidation by CELLOX1 occurs at the glucose residue at the reducing ends of TRIA and TETC (Figure 5), indicating that only molecules that have β-(1,4)-linked glucose at the reducing end are suitable substrates of the enzyme (Figure S4). In summary, CELLOX1 and CELLOX2 are characterized by comparable enzymatic properties in terms of both substrate specificity and pH-dependent activity, and act on CDs and MLGs with a β-1,4-glucose but not a β-1,3-glucose at their reducing ends.

**Figure 4.**
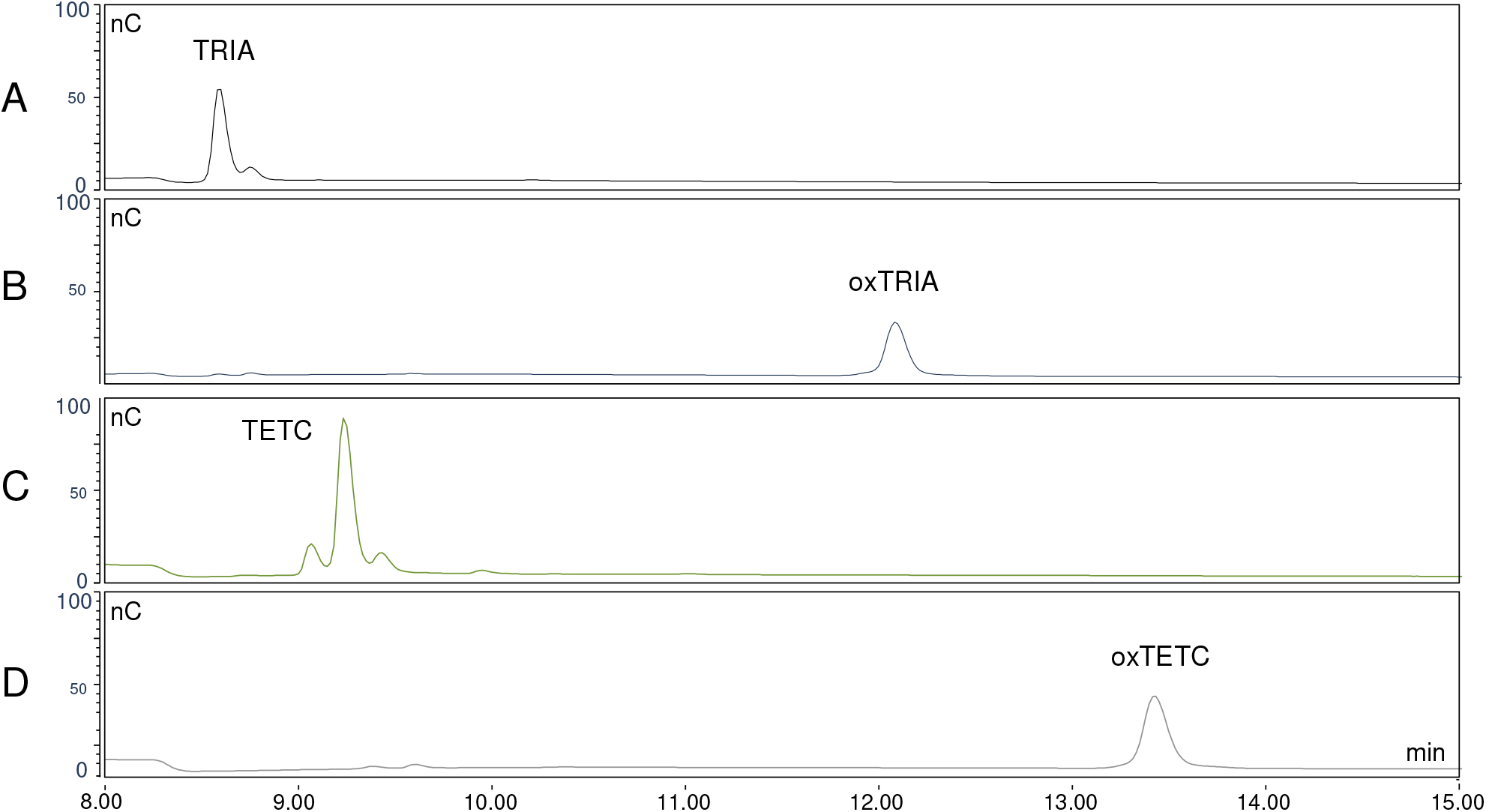
TRIA and TETC are substrates of CELLOX1. HPAEC-PAD of (A) native and (B) CELLOX1-treated TRIA, and of (C) native and (D) CELLOX1-treated TETC.

**Figure 5.**
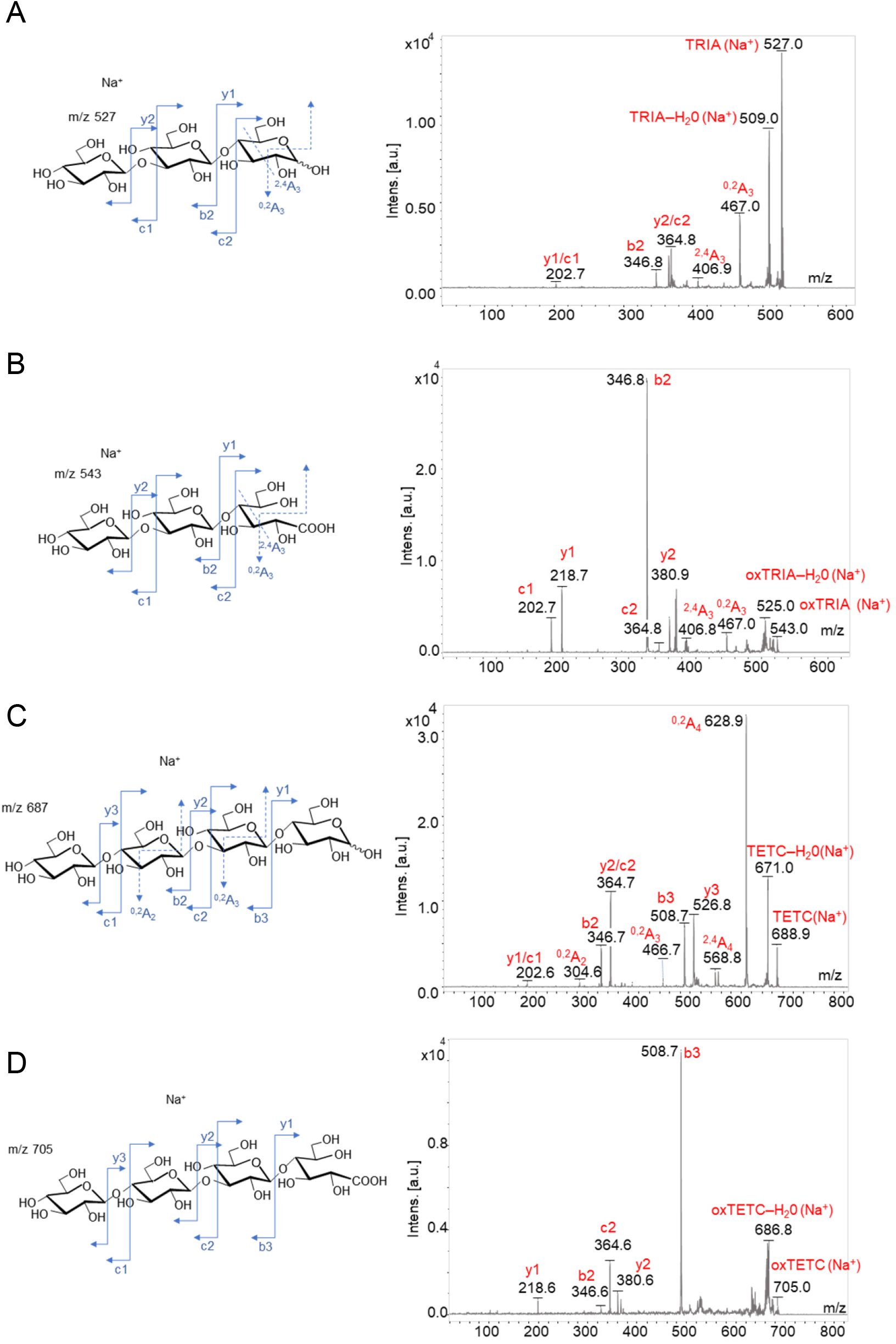
CELLOX1 oxidizes the glucose residue at the reducing ends of TRIA and TETC. MS analyses of (A) native and (B) CELLOX1-treated TRIA, and of (C) native and (D) CELLOX1-treated TETC. For each oligosaccharide, the respective structure (left) and mass spectra (right) of positive ions determined by MALDI-TOF/TOF analysis are shown. The fragment ion peaks are labelled according to the nomenclature proposed by Domon and Costello (1988). [solid line: glycosidic linkage fragments, dashed line: cross-ring fragments].

Since the DAMP capacity of molecules such as OGs and CDs is substantially impeded by oxidation (Benedetti et al., 2018; Locci et al., 2019; Pontiggia et al., 2020), we investigated whether the elicitor activity of MLGs is also impaired by the enzymatic oxidation, by assaying native and oxidized MLG preparations on Arabidopsis Columbia-0 (Col-0) seedlings. The genes *WRKY53, FRK1* and *CELLOX1* were analyzed as readouts of defenses induced by MLGs and CDs (Barghahn et al., 2021; Rebaque et al., 2021). TRIA and TETC and the oxidized counterparts oxTRIA and oxTETC were tested in parallel with OGs and CD3, as positive controls of defense gene induction. As shown in Figure 6, TRIA and TETC induced the expression of all the three genes analyzed, while oxTRIA and oxTETC exhibited a strongly reduced elicitor capability.

**Figure 6.**
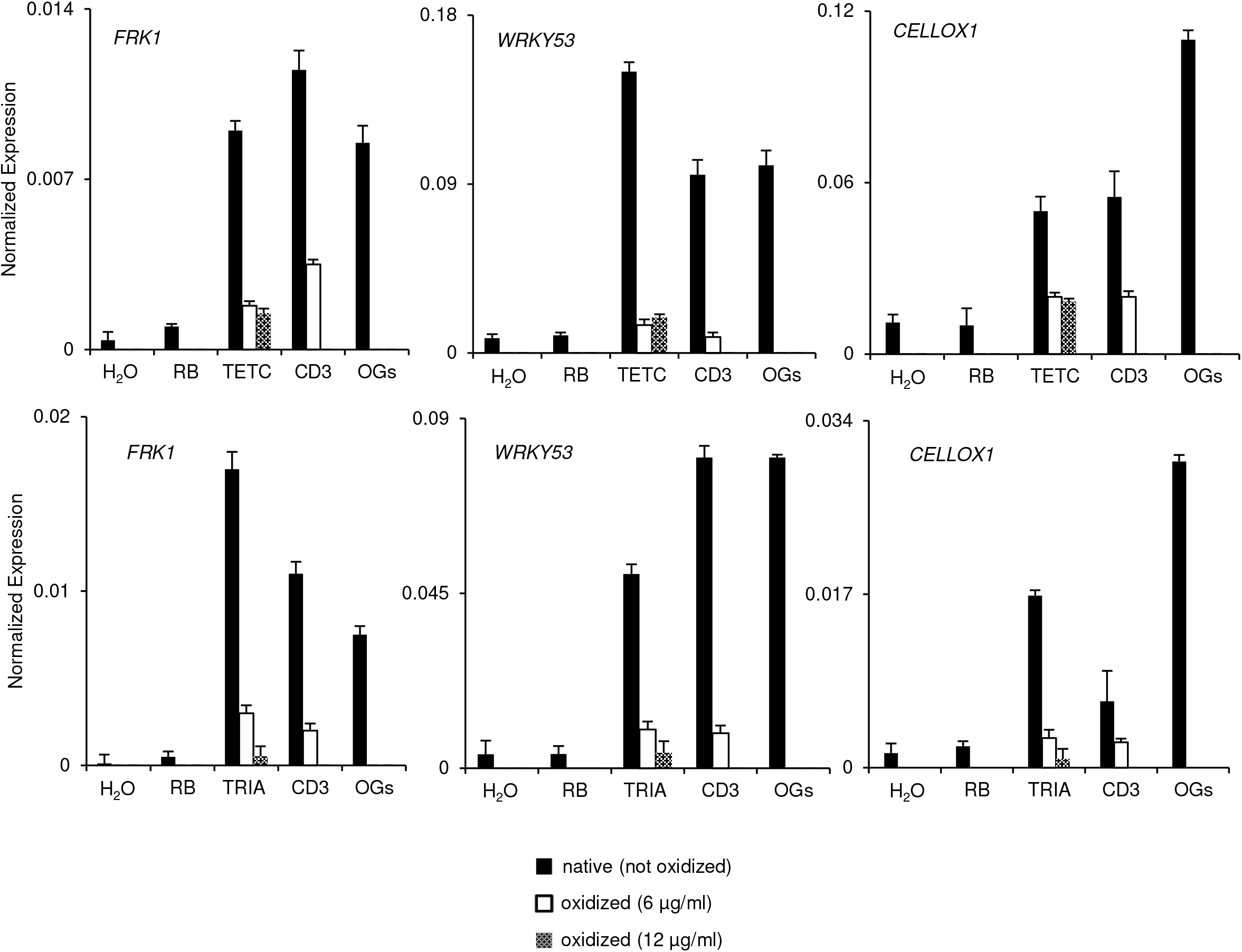
The enzymatic oxidation of TETC and TRIA impairs their elicitor activity. Col-0 wildtype seedlings were treated for 30 min with native and oxidized TETC (upper panels) and TRIA (lower panels). Native TETC and TRIA, and native and oxidized CD3 were used at a concentration of 6 μg/ml. Oxidized TETC and TRIA were used also at a concentration of 12 μg/ml. Treatment with water (H_2_O) and reaction buffer (RB) was used to evaluate the basal gene expression, whereas treatment with oligogalacturonides (OGs; 25 μg/ml) was as a positive control of defense gene induction. Transcript levels of *FRK1, WRKY53* and *CELLOX1* were analyzed by quantitative RT-PCR and normalized to *UBQ5* expression.

### CELLOX2, unlike CELLOX1, is not involved in immunity against *B. cinerea*

To assess whether *CELLOX2* is involved in immunity, commercially available seeds carrying T-DNA insertions within the coding sequence of *cellox2* in the Col-2 background as well as *cellox1* mutants in the Col-0 background were analyzed. As determined by semiquantitative RT-PCR analysis (sqRT-PCR), *cellox1* and *cellox2* are both knockout mutants (Figure S5). The two mutants were infected in parallel with *B. cinerea* and the area of the lesions was measured 48 h after inoculation (hpi). The average lesion area of the *cellox1* knockout mutant was significantly smaller than that of the Col-0 wild type (Figure 7). The lesion area was reverted to wild type size in *cellox1* plants complemented with a pCELLOX1::*CELLOX1* (wild type) cassette *(cellox1-c*), which had levels of *CELLOX1* transcripts similar to those of Col-0 plants (Figure S6). Instead, the average lesion area in the *cellox2* mutant was not significantly different from that of the wild type, Col-2, indicating that *CELLOX2* is not involved in defense against *B. cinerea* (Figure 7). The lack of involvement of *CELLOX2* in immunity against *B. cinerea* supports *in-silico* data showing that *CELLOX2*, unlike *CELLOX1*, is not induced during the immune response (Figure S7). Indeed, the expression of *CELLOX2* in Col-0 seedlings upon treatment with the MAMPs flg22 and elf18, or the DAMPs OGs and CD3 was undetectable, confirming that the gene is neither expressed in the vegetative parts of the seedling nor upon induction with elicitors (Figure S8).

**Figure 7.**
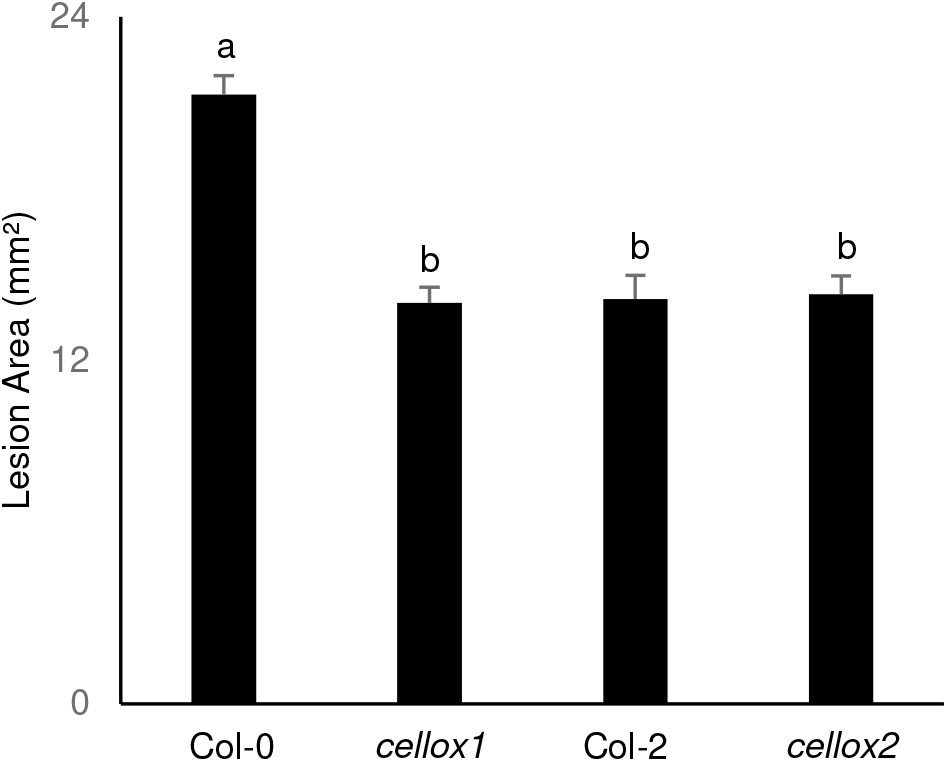
Infection of CELLOX null mutants with *Botrytis cinerea.* Lesion area (mm^2^, average ± SE; four biological replicates) in leaves of of transgenic *Arabidopsis cellox1* and *cellox2* mutants 48 h after inoculation with *Botrytis cinerea* spores. Col-0 was used as control of *cellox1;* Col-2 was used as control of *cellox2.* ANOVA followed by post-hoc Tukey’s HSD test (P < 0.05).

### CELLOX1 and CELLOX2 influence the seed coat structure and mucilage porosity but not their carbohydrate composition

*In silico* analysis of the spatial distribution of *CELLOX1* and *CELLOX2* shows that both genes are expressed in the seed coat during development, although with a different temporal profile (Figure S9). *CELLOX2* is expressed at the pre-globular and globular stages with a higher level at the heart stage and is not expressed at the later stages (linear cotyledon and maturation green), when *CELLOX1* is instead expressed. On the other hand, only *CELLOX2* is expressed in the embryo, at the maturation green stage and in the peripheral endosperm (Figure S9). In accordance with the *in-silico* analysis, *CELLOX1* and *CELLOX2* transcripts were detected in the seeds of the wild type mutants and were absent in the seeds from the T-DNA insertional mutants *cellox1* and *cellox2*, respectively (Figure S10).

Coat and mucilage are both formed by the seed coat epidermis after cell expansion by deposition of secondary wall. The mucilage can be distinguished in non-adherent mucilage, which easily dissociates from the seed and is mainly composed by the pectin RG-I, and the coat-adherent mucilage, composed of other types of pectin polysaccharides, cellulose and hemicellulose. Structures are present in the adherent mucilage, i.e., the *columella*, and rays that extend perpendicularly above the columella. Both structures are stained by cellulose-specific dyes, which also stain a more diffuse area between the rays (Griffiths and North, 2017; Di Marzo et al., 2022). Scanning electron microscopy (SEM) analysis of dry seeds (Figure 8A) revealed that the *columella* area was significantly smaller in *cellox1* but not in *cellox2*, compared to the wild types, Col-0 and Col-2, respectively (Figure 8B), whereas the radial cell walls of coat epidermal cells of both *cellox1* and *cellox2* were thicker (Figure 8C). No significant differences were detected in the area of dry and hydrated seeds of the *cellox1* and *cellox2* mutants compared to the respective wild types (Figure 8D).

**Figure 8.**
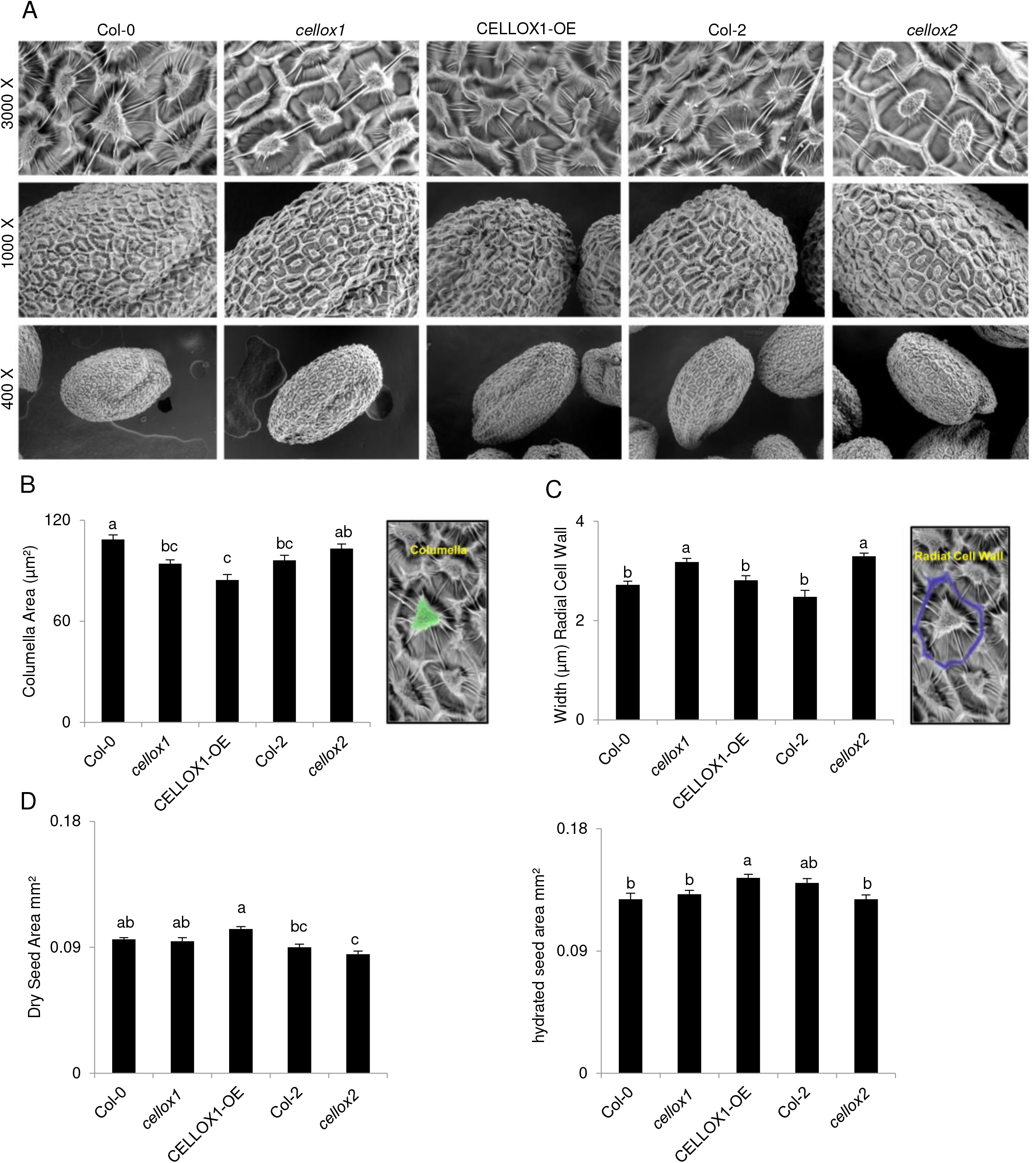
Scanning electron microscopy of seeds with altered CELLOX expression. A) In the upper panels, close up view of epidermal cells for each mutant (3000 x). In the lower panels it is possible to observe different magnifications at 1000 x and 400 x, which show the entire shape of each mutant seed. B) area (μm^2^) of seed columella in the indicated genotypes, highlighted in green in the picture on the right. C) Width of radial cell wall (μm) for each mutant. Radial cell wall are highlighted in blue in the picture on the right. D) Area of dry seed (left) and hydrated seed (excluding mucilage, right) for each mutant. Col-0 and Col-2 were used as controls for *cellox1* and *cellox2* mutants respectively. Different letters indicate statistically significant differences according to ANOVA followed by post-hoc Tukey’s HSD test (P < 0.05). Columella area and width of radial cell wall were measured using Image J software.

Seeds were hydrated and stained with Ruthenium Red (RR), which stains pectin, and fluorescein isothiocyanate (FITC)-dextran 70, which is used to examine the mucilage porosity because it does not normally penetrate the mucilage adherent layer and is partially excluded from the cellulosic rays (Willats et al., 2001; Voiniciuc et al., 2015). No significant difference was observed in the RR-stained halo of the seeds in the *CELLOX1* null and overexpressing plants, whereas a reduced halo was visible in the *cellox2* mutant (Figure 9A). Confocal microscopy analysis of the FITC-dextran-stained seeds revealed a dense and compact adherent mucilage layer in Col-0 and Col-2, and a less dense layer in both the *cellox1* and *cellox2* mutants, where the dye penetrated through the mucilage and between the cellulose rays and reached the most inner part of the seed coat surface (Figure 9B). The seed features of the wild type and mutant seeds are summarized in table 1. Taken together, these results suggest a role of both CELLOX1 and CELLOX2 in determining the porosity of the seed adherent mucilage, with CELLOX2 likely also contributing to the formation of the mucilage layer. Seed morphology analyses was extended to the seeds of the available plants overexpressing CELLOX1 (CELLOX1-OE) (Locci et al., 2019). These also showed alterations of the seeds (Table 1): an increased size of both dry and hydrated seeds, a decreased *columella* area (Figure 8B) with normal radial wall (Figure 8D), and a lower density of adherent mucilage (Figure 9B). Neither the null *cellox* nor the overexpressing seed showed significant defects in germination (Figure S13). Overall, these results point to a role of both CELLOX1 and CELLOX2 in the formation of radial cell walls and in the regulation of mucilage porosity, and to a more important role of CELLOX1 in *columella* formation. Both enzymes do not play a major role in germination.

**Figure 9.**
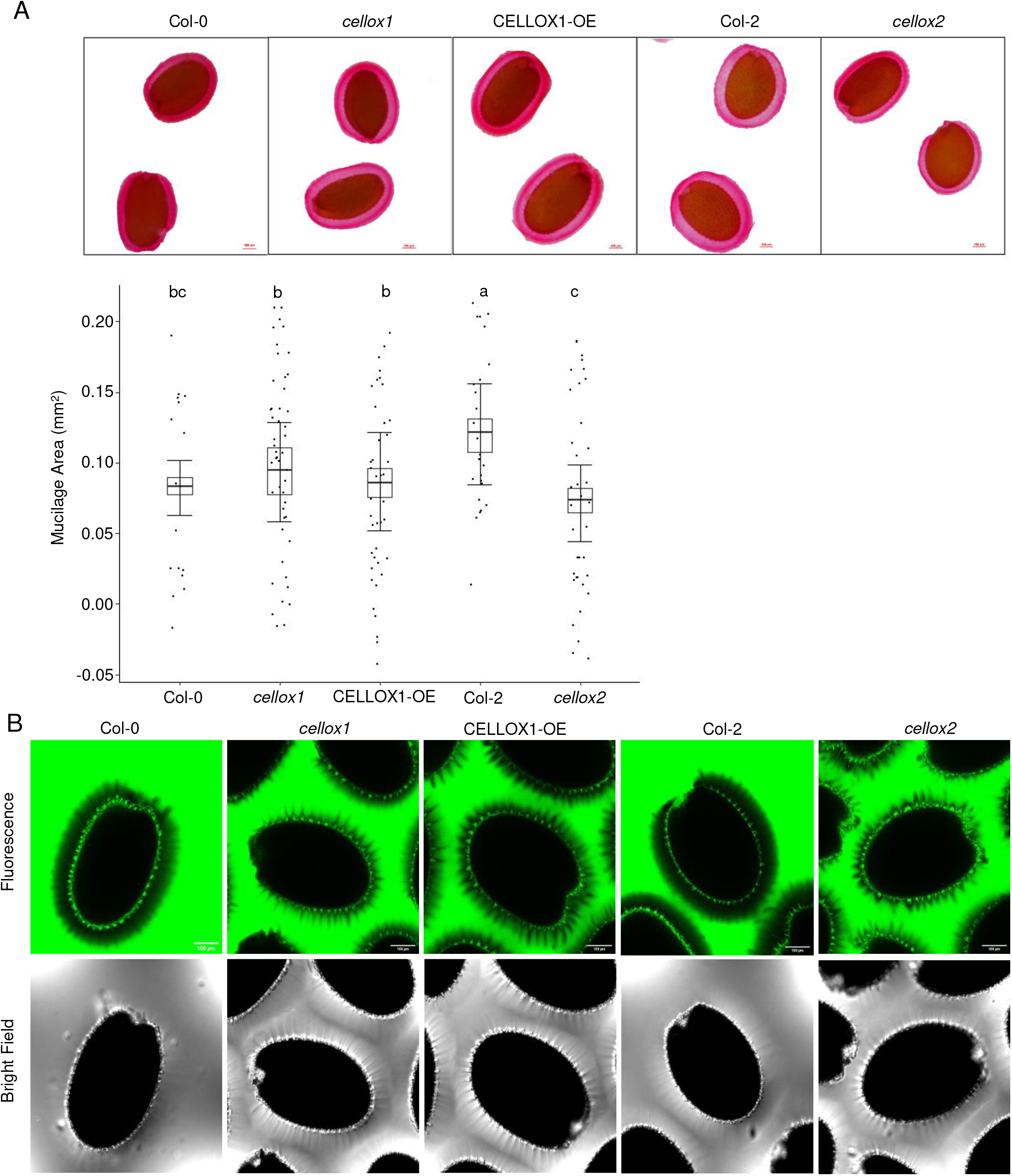
Mucilage analysis in *CELLOX* mutants. **A)** (Top panel) Adherent seed mucilage, detected by Ruthenium Red (RR) staining, which highlights pectin. Images were taken at optical microscope (4 X, scale bar 100 μm). Ecotypes Col-0 and Col-2 were used as controls for *cellox1* and *cellox2*, respectively. (Bottom panel) Box plot of adherent mucilage area, measured after hydration and RR staining. Average ± SE of at least three replicates; different letters indicate statistically significant differences according to ANOVA followed by post-hocTukey’s HSD test (P < 0.05). B) Mucilage density, analyzed by staining with fluorescein isothiocyanate-labelled dextran ((FITC-Dextran 70), which does not normally penetrate the mucilage adherent layer and is partially excluded from the cellulosic rays (Voiciniuc et al., 2015). Pictures show that the fluorophore is completely excluded from the adherent mucilage layer of Col-0 and Col-2 (dark halo), indicating a dense layer, and from the rays. Instead, it penetrates well the mucilage of *cellox1* and *cellox2* mutants, and partially the mucilage of CELLOX1-OE. Bright filed images show that thin rays are visible only in the null and the overexpressing plants, likely because a less dense mucilage “unmasks” the rays, the structure of which however appears not to be affected. Scale bar 100 μm. Confocal images.

**Table 1.** Phenotypic features^a^ observed in CELLOX mutant seeds.

|  | *cellox1* | *cellox2* | **CELLOX1-OE** |
| --- | --- | --- | --- |
| **Dry seed area** | = | = | ↑ |
| **Hydrated seed area** | = | = | ↑ |
| **Columella area** | ↓ | = | ↓ |
| **Width of radial cell walls** | ↑ | ↑ | = |
| **Thickness of the adherent mucilage layer** | = | ↓ | = |
| **Porosity of adherent mucilage layer** | ↓ | ↓ | ↓ |

<sup>a</sup> Symbols indicate a similar (=), increased (↑) and decreased (↓) phenotype compared to the respective wild type.

The observed alterations prompted us to investigate the monosaccharide composition of both the mucilage and the seed coat of the *cellox* mutants and the CELLOX1-OE plants. After extraction of the mucilage fraction, an Alcohol Insoluble Solids fraction was obtained from the de-mucilaged seeds (dsAIS). Both fractions were hydrolyzed with TFA (Figure S11), which solubilizes matrix components, and the residue of the TFA-hydrolyzed dsAIS was also hydrolyzed with H2SO4, to solubilize cellulose (Figure S12). HPAEC-PAD chromatography analysis showed that the total amount of sugars did not show major changes among the *cellox*, CELLOX1-OE, Col-0 and Col-2 genotypes in all the hydrolysates (Figure S11, Figure S12). In agreement with literature data, Rha and GalA were the most abundant monosaccharides in the mucilage, whereas Ara and Gal were the major components in the TFA-hydrolyzed dsAIS (Figure S11). In the dsAIS samples obtained upon sequential TFA-H2SO4 hydrolysis, Glc (~56 %) was the major component, with lower amounts of Ara (~14 %), Gal (~14 %), Xyl (~12 %) and traces of Rha. In conclusion no substantial differences between the carbohydrate compositions in the different genotypes were evident within the limitations of the utilized procedures.

## DISCUSSION

The member of the BBE-l protein family encoded by the gene *CELLOX2/BBE23/At5g44360* has been characterized in this work. CELLOX2 shares 78.88 % amino acid identity and several structural features with the previously characterized CELLOX1 (Locci et al., 2019). A type IV active site is present in both CELLOX1 and CELLOX2; this type of active site is found in BBE-l proteins from fungi, lower plants (Daniel et al., 2017) and monocotyledons (Messenlehner et al., 2021). CELLOX2 also shares with CELLOX1 enzymatic features such as the capability of oxidizing β-1,4-glucan oligomers and a marked alkaline dependent activity. Notably, both CELLOX1 and CELLOX2, oxidize the MLGs, recently identified as DAMPs capable of activating the immune responses in Arabidopsis, rice and barley (Barghahn et al., 2021; Rebaque et al., 2021; Yang et al., 2021). MLGs are structural components of the cell wall of some monocots such as the *Poaceae*, and may also be present in algae, bryophytes and different *Equisetum* species (Sørensen et al., 2008; Salmeán et al., 2017). On the other hand, although they induce immunity in Arabidopsis, MLGs are not present in the cell wall of dicots (Rebaque et al., 2021), and therefore, since their presence has been reported also in fungi, oomycetes and bacteria (Pettolino et al., 2009; Samar et al., 2015), their action on dicots is more related to their role as MAMPs. The oxidizing activity of CELLOXs on endogenous and exogenous cell wall-derived oligosaccharides acting either as DAMPs or MAMPs reinforce the view that members of the BBE-l family are devoted to the regulation and homeostasis of the elicitors of plant defense, whose accumulation may cause hyper-immunity (De Lorenzo et al., 2018; Pontiggia et al., 2020; De Lorenzo and Cervone, 2022).

Notably, both CELLOXs and the receptor kinases IMPAIRED IN GLYCAN PERCEPTION 1 and 4 (IGP1 and IGP4) recognize as substrates and ligands, respectively, cellulose fragments but not MLG that has a β-(1,3)-linked glucose at the reducing end (Martín-Dacal et al., 2022). IGP1 is also known as the Cello-Oligomer Receptor Kinase 1 (CORK1), required for CD3 perception and PTI activation in Arabidopsis (Tseng et al., 2022). IGP1/CORK1 and its paralogs IGP2/3 and EGP4 have Leucine-Rich-Repeat and Malectin domains in their ectodomain (LRR-MAL), and loss-of-function mutations in these proteins affect PTI activation mediated by MLG43 and CD3, but not by the chitin fragment Chi6, which also is not a substrate of CELLOXs (see Figure 3B). These observations suggest that CELLOXs and LRR-MAL receptor kinases share structural requisites for specific binding of CD fragments. Whether IGPs bind MLGs that have a β-(1,4)-linked glucose at the reducing end is not known yet.

The inactivating effect of CELLOX1 on the elicitor activity of CDs has an impact on the outcome of the infection against *Botrytis cinerea*, since *cellox1* mutants are more resistant to this fungus. On the other hand, plants overexpressing CELLOX1 are also more resistant to *B. cinerea* and this has been explained with a less efficient utilization of oxidized oligosaccharides as a carbon source by the fungus (Locci et al., 2019). The dual impact of these enzymes on the infection process by *B. cinerea* may grant resistance even when the homeostatic function of CELLOXs lowers the levels of elicitor active molecules, which otherwise may trigger deleterious effects of hyper-immunity. Whether and how the production of H_2_O_2_ by the enzyme contributes to enhance the defense responses is still to be investigated.

*CELLOX2*, unlike *CELLOX1*, was found in this paper not to be involved in immunity of seedlings against *B. cinerea* in agreement with the lack of induction of its expression during immunity, as indicated by available transcriptome data. This observation prompts to further investigate possible roles of this gene in processes not related to immunity. Notably CW-derived DAMPs such as OGs and CDs have often been reported to work as signals at the crossroads between immunity and development (De Lorenzo and Cervone, 2022). *CELLOX1* and *CELLOX2* show different expression profiles in the Arabidopsis tissues. *CELLOX2* is not expressed in seedlings or in adult plants but only in the seed coat during the early intermediate stages of seed development. Also *CELLOX1* is expressed in the seed coat, but only when *CELLOX2* shows a decreased expression, suggesting a concerted regulation, the biological significance of which is not yet clear. The seed coat surrounds the embryo throughout its development and dormancy, providing protection from mechanical damage and pathogens and regulating germination, dispersal and seed viability in hostile environments (Huang et al., 2008; Ben-Tov et al., 2015). Other members of the BBE-l family do not appear to be expressed in the seed coat; thus, *CELLOX2* and *CELLOX1* may play a unique role in this tissue as oxidases. Indeed, an altered CELLOX expression alters the seed structure (see Table 1). Both null and overexpressing CELLOX mutants show defects of the seed coat and mucilage. *Columella* and rays, mainly cellulosic, are secondary wall depositions formed by the seed coat epidermis, after deposition of mucilage, mainly composed of the pectin RG-I (Griffiths and North, 2017). Since MLGs are not present in Arabidopsis, and sugar composition of the seed coat is not altered, the observed defects suggest that the structure of cellulose, rather than its abundance, is mainly affected by CELLOX. The effects of an altered CELLOX function in plants including those that possess MLGs in the cell wall such as *Poaceae*, remains to be investigated. It is worth noting that growth defects have been observed in both MLG over-accumulating or MLG-deficient plants (Burton et al., 2011; Kim et al., 2018; Bain et al., 2020).

Interestingly, the mutants defective in *CELLULOSE SYNTHASE* 5 *(CESA5)* show a reduced *columella* area and a reduced and less dense adherent mucilage layer, as observed in the null *cellox* mutants (Sullivan et al., 2011). These results suggest that that the structure of cellulose plays an important role in the adhesion of pectin and therefore of mucilage in the seed (Griffiths et al., 2014; Griffiths and North, 2017). However, important differences are displayed by the two types of mutants. Seed size is reduced in the *CESA5* mutant, whereas it is unaffected in *cellox* mutants (Mendu et al., 2011). Moreover, triple mutants defective in *CESA5, CESA2* and *CESA9* show a reduction in the radial walls, which instead are thicker in both *cellox* mutants.

While the germination is not significantly affected in *cellox* mutants, it is not known whether a homeostatic control of defense signaling is played by CELLOXs and is needed during seed development. Defense molecules, including chemical metabolites such as hydroxyphenols, and hydrogen cyanide released upon hydration, pathogenesis-related proteins such as β-1,3 glucanases, nucleases, proteases and chitinases, are often located in the seeds (Raviv et al., 2017). It is worth mentioning that mutants associated with defense against biotic stresses may also display an altered seed phenotype. For example, loss-of-function mutants in TRANSPARENT TESTA GLABRA2 (TTG2), a WRKY transcription factor involved in responses to pathogen attack and mechanical stresses, show absence of internal domes of *columellae* and overlying mucilage. A different spatial and temporal regulation of the expression of two BBE-l proteins with similar enzymatic activity may allow their action as sentinels of danger in different organs and tissues against several pathogens or/and during different events of plant development, also by controlling the entity of the immune response and producing signal molecules such as H_2_O_2_ or causing localized alterations of the cell wall structure. In addition, both CELLOXs and OGOXs may provide H_2_O_2_ necessary for the plant peroxidase-catalyzed oxidation of substrates such as auxin and monolignols in the apoplast (Scortica et al., 2022).

In conclusion this paper reports that CELLOX1 and CELLOX2 act as DAMP/MAMP oxidases displaying oxidative activity not only on CDs but also on MLGs with a different pattern of expression. While CELLOX1 is involved in immunity against *B. cinerea*, both CELLOX1 and CELLOX2 influence seed coat structure and mucilage porosity confirming that BBE-l proteins deserve deeper insights and further studies both in the field of immunity and development. The function of many BBE-l proteins is still unclear, and many members of this family may turn out to be enzymes whose substrates are cell wall poly- and oligosaccharides.

## MATERIALS AND METHODS

### Plant material and growth

Arabidopsis *s*eeds were sterilized (Locci et al., 2019), and grown on MS/2 medium at 22 °C and 70 % relative humidity under a 16 h light/ 8 h dark cycle. Adult plants were transferred on soil at 22 °C and 70 % relative humidity under a 16 h light/8 h dark cycle (approximately 120 μmol m-2 s-1). Plants used for infections with *Botrytis cinerea* have been grown under a 12 h light/ 12 h dark cycle at 22 °C.

T-DNA insertional lines were obtained from the European Arabidopsis Stock Center (NASC). For *At4g20860 (CELLOX1)* and *At5g44360 (CELLOX2)* the lines SALK_131268 (NASC ID N631268), here indicated as *cellox1*, and WiscDsLox472A5 (NASC ID N857391; *cellox2)*, respectively, were genotyped. Overexpressing lines of *CELLOX1* (CELLOX1-OE, line #9.7) (Locci et al., 2019) and *OGOX1* (Benedetti et al., 2018) were also used.

*N. tabacum* was used for transient expression of GFP-tagged CELLOX2 *(At5g44360.1/BBE23)* in agro-infiltrated leaves for localization analysis by confocal microscopy.

### Preparation of constructs for the expression of tagged BBE23

Signal peptide prediction of BBE23 was performed (Data S1) using the online tool SignalP-5.0 (https://services.healthtech.dtu.dk/service.php?SignalP-5.0). Then, the sequence of mature BBE23 protein (UniprotKB: Q9FKV2, aa 23-532) was reverse-translated into a codon-optimized sequence in accordance with the codon usage of *P. pastoris* by using the software OPTIMIZER (http://genomes.urv.es/OPTIMIZER/) (Puigbo et al., 2007) and fused downstream of the Flag-His-SUMOstar sequence (FSH, LifeSensor Inc.; https://lifesensors.com/wp-content/uploads/2019/09/2160_2161_Pichia_SUMOstar_Manual-1.pdf), which includes a Flag epitope (DYKDDDDK) and 6x histidine (6xHis; HHHHHH) tag sequence. The His-tagged BBE23 variant (H-BBE23) was obtained in accordance with the same protein design of H-OGOX1 (Scortica et al., 2021). Both the FHS-BBE23 and H-BBE23 codon-optimized fusion genes were synthesized by GeneArt (Life Technologies) and cloned in pPICZαB expression vector (Invitrogen, San Diego, CA).

The Gateway system vectors were used to obtain the constructs *35S::CELLOX2-GFP* for *Agrobacterium tumefaciens*-mediated transient expression in leaves. pDONRTM221 (Invitrogen) Kanamycin and Zeocin resistant, and the pK7FWG2 (V141) for the eGFP C-Term tag and the pB7RWG2 (V215) for the mRFPC-Term tag, kanamycin resistant (Karimi et al., 2002).

### Heterologous expression of BBE enzymes in *P. pastoris*

H-BBE23 and FHS-BBE23 were expressed in *P. pastoris* as previously described (Scortica et al., 2021; Scortica et al., 2022). In order to facilitate the detection of BBE-l proteins, the culture filtrates from methanol-induced cultures were treated with PNGaseF (New England Biolabs), and then evaluated by immune-decoration analysis using a monoclonal His-antibody (AbHis, Bio-rad). Pichia transformants expressing FHS-CELLOX was already available (Scortica et al., 2022).

### Purification of FHS-CELLOX and FHS-BBE23

The purification of FHS-CELLOX and FHS-BBE23 was prepared as previously described (Scortica et al., 2022). Purity of the protein preparation was assessed by SDS-PAGE/Coomassie blue staining whereas the concentration of FHS-BBE23 was measured by UV absorbance (ε280nm: 88030 M^-1^ cm^-1^).

### Biochemical characterization of FHS-CELLOX and FHS-BBE23

The xylenol-orange assay was initially used to analyze the enzyme activity of pure FHS-CELLOX and FHS-BBE23 (Benedetti et al., 2018). The enzymes were used at a concentration of 150 ng mL^-1^ (5 nM) and the reaction temperature set to 25°C. Analysis of substrate specificity was performed in 50 mM Tris-HCl, pH 7.5, and 50 mM NaCl by using the following substrates (0.4 mM each): monosaccharides (D-glucose, D-xylose, D-galactose), disaccharides (D-cellobiose, D-lactose, D-maltose), cellotriose (CD3), cellotetraose (CD4), cellohexaose (CD6), chitotriose (Chi3), chitohexaose (Chi6), 3-β-D-cellotriosyl-glucose (TETB), 3-β-D-glucosyl-cellobiose (TRIA), 3,1-β-D-cellobiosyl-glucose (TRIB), oligogalacturonides (OGs), N-acetyl-glucosamine (NAG), and a mixture of 3-β-D-cellobiosyl-cellobiose and 3-β-D-glucosyl-cellotriose (TETC). All monosaccharides, disaccharides and CDs were purchased from Sigma-Aldritch (St. Louis, USA) whereas MLGs (TETB, TRIA, TRIB and TETC) and chitin oligosaccharides were purchased from Megazyme (Bray, Ireland). OGs were prepared as previously described (Pontiggia et al., 2015) The μmoles of H_2_O_2_ generated by each BBE-l protein towards different substrates were evaluated at three different time-points from the linear phase of each reaction and then averaged. Relative activity (%) was calculated by applying the following formula: [(μmol H_2_O_2_ *vs* substrate “X”).(μmol H_2_O_2_ *vs* optimal substrate)^-1^ ×100%]. Buffers at different pH values ranging from 5 to 12 were supplied with cellobiose (0.4 mM) and used for evaluating the pH optimum of the enzymes. The μmoles of H_2_O_2_ generated by each BBE-l protein in the pH range 5-12 were evaluated at three different time-points from the linear phase of each reaction and the relative activity (%) was calculated at each pH value as [(μmol H_2_O_2_ at pH_x_).(μmol H_2_O_2_ at pH_opt_)^-1^ ×100%] in order to obtain the pH-dependent activity curve.

### Preparation of native and oxidized oligosaccharides for elicitation assays

Commercial TRIA, TETC and CD3 were separately dissolved in Reaction Buffer [RB; 50 mM Tris-HCl, pH 8.8, 50 mM NaCl, bovine liver catalase (1 μg mL^-1^, Sigma-Aldrich)] and incubated at 28°C for 24 h in the presence and in the absence of purified FHS-CELLOX1 (20 μg). After incubation, a small aliquot from each mixture was analyzed by HPAEC-PAD and xylenol-orange assay to confirm, respectively, the complete oxidation of the vary oligosaccharides and the absence of residual H_2_O_2_. Enzymes were denatured at 65 °C for 20 min, and samples were filtered by using a Vivaspin Turbo4 Ultrafiltration Unit (10.000 MWCO PES; Sartorius, Gottinga, Germany) to remove the proteins.

### Analysis of elicitor activity of oxidized MLGs

To evaluate the elicitor activity, 12-day-old Col-0 seedlings were grown in liquid MS/2 medium and treated for 30 min with the ultrafiltered preparations of native and oxidized TRIA, TETC, and CD3. OGs (25 μg mL^-1^) were also used as positive controls. Col-0 seedlings treated with ultrapure sterile water (H_2_O) and ultrafiltered RB were used to evaluate the basal gene expression.

### Chromatographic and mass spectrometric analysis of standard and modified oligosaccharides

Oligosaccharides were analyzed by HPAEC-PAD with an ICS-3000 apparatus (Dionex Corporation) according to Pontiggia et al. (2015) with modifications. The sample (10 μl) was injected on a CarboPac PA200 3×250 mm analytical column with a guard column (Thermo Fisher) kept at 35 °C. The flow rate was 0.4 ml min^-1^ and eluents A (0.05 M NaOH) and B (1 M Na-acetate in 0.05 M NaOH) were applied as follows after injection: a 5min linear gradient to 5 % B and then 8 % of B in 9 min, followed by a 2-min linear gradient to 12% B a 1-min gradient to 50 % B. 100 % B was kept for 5 min before returning to 100 % A and equilibrating for 10 min.

MALDI-MS analysis was performed according to Scafati et al. (2022) with modifications. MS spectra of the sodium adducts of oligosaccharides [M + Na]^+^ were performed with an UltraFlex MALDI-TOF/TOF mass spectrometer (Bruker, Germany). In the positive ionization mode, the MS2 on standard and modified MLGs produced cleavage fragments, generating B-, A-, C- and Y-ions according to the nomenclature proposed by Domon and Costello (1988).

### RNA extraction and gene expression analysis

Total RNA was extracted from Arabidopsis seedling tissues with NucleoZOL Reagent (Macherey-Nagel) according to the manufacturer’s protocol and gene expression analysis was carried out as previously described (Locci et al., 2019).

### Infection assay of *Botrytis cinerea* on Arabidopsis leaves

*B. cinerea* was grown and infections performed as previously described (Locci et al., 2019). Lesion size was determined by measuring the necrotic area (mm^2^) by using ImageJ software.

### Agroinfiltration

The infiltration buffer [50 mM MES, pH 5.6 (pH adjusted with KOH), 2 mM sodium phosphate buffer, pH 7, 200 μM acetosyringone, 0.5 % glucose] was used to resuspend the transformed *Agrobacterium tumefaciens* for inoculation. OD at 600 nm was measured to have a final value of 0.1 and 0.2 for each sample for infiltration of *N. tabacum* leaves using needless a syringe. The infiltrated leaves were left for 48 h in the greenhouse before confocal microscope visualization, using a Zeiss LSM 780 microscope.

### Treatments for the analysis of *CELLOX2* expression

For the analysis of the expression of *CELLOX2* by semiquantitative RT-PCR (sqRT-PCR), 10-day-old Col-0 seedlings grown in liquid medium were treated 1 h with the elicitors flg22 (10 nM), elf18 (10 nM), OGs (50 μg mL^-1^), and cellotriose (CD3; 25 μg mL^-1^).

### RNA extraction from dry seeds for transcript analysis

RNA extraction from dry seeds of Arabidopsis was performed according to Zhao et al. (2019) with modifications. Dry seeds (30 mg) were frozen in liquid nitrogen and grounded at mixer miller. Extraction buffer (0.4 M LiCl, 0.2 M Tris (pH 8.0), 25 mM EDTA, 1 % SDS) and chloroform were added to the grounded seeds and mixed. Then, centrifugation followed 16000 x g for 3 min. The upper phase was transferred to a new 1.5 mL Eppendorf tube and a similar amount of phenol (1:1) was added and vortexed. After 5 min of incubation in ice, chloroform was added in an amount corresponding to the 2 % of the total volume. 3 minutes of centrifugation at 16000 x g followed. The upper phase was collected and transferred to a new 1.5 mL tube containing 8 M LiCl (1/3 total volume), and mixed. The solution was left for 30 minutes at – 20 °C and centrifuged 16000 x g for 15 minutes at 4 °C. The supernatant was discarded. Washing in ethanol followed, sequentially 75 % EtOH and 100 % EtOH, supernatant was discarded, and the pellet was left to dry for few minutes. Sterile H_2_O nuclease free water was added. Gene expression analysis was performed as previously indicated.

### Scanning Electron Microscopy (SEM)

SEM was used to examine epidermal morphology of dry seeds. Wild-type and *cellox* mutants seeds were mounted without any preparation on aluminum stubs and observed by Scanning Electron Microscopy (FESEM Gemini 500, Zeiss) equipped whit Peltier MK3 Coolstage (Zeiss) at 10 kV, 20Pa (VP), 4°C. The SEM observations were carried out at different magnifications (400 X, 1000 X and 3000 X).

### Ruthenium Red Staining and seed and mucilage areas measurement

Mucilage area was measured according to Wang et al. (2019), with some modifications. About 50 seeds from batches grown simultaneously and under same conditions were used to quantify the seeds and adherent mucilage areas. Seeds were placed in a clean (but not necessarily sterile) microfuge tube. Distilled H_2_O was added for seeds imbibition for the seeds to hydrate and release the mucilage from the epidermal seed coat, and the tube was shaken (~ 300 rpm) on an orbital shaker for 40’ at room temperature to remove the non-adherent layer. Supernatant was carefully removed and replaced with 0.01 % ruthenium red solution for 5’. The Ruthenium Red solution was removed and replaced with distilled H_2_O for washing before the observation and the acquisition of images at optical microscope (4 X) with a bright field. The area (mm^2^) of the whole seed and adherent mucilage was measured using the software ImageJ (Schneider et al., 2012) and data were normalized on the average Col-0 seed area. The experiment was repeated three times.

### FITC dextran staining

Seed mucilage density was evaluated by using Fluorescein isothiocyanate (FITC)-dextran staining [FITC-Dextran 70 (Sigma-Aldrich; 46945) (Voiniciuc et al., 2015)]. The seeds were soaked in 100 mM sodium acetate, pH 4.5, for 10 min before adding FITC–Dextran 70 (1 mg mL^-1^) and then incubated for 30 min on a shaking platform (125 rpm) in dark condition. Stained seeds with FITC were detected with a Leica TCS SP5 II confocal system (488 nm of excitation and 500-540 nm of emission).

### Extraction of total mucilage

Total mucilage extraction was performed as described by Voiniciuc et al. (2015) with some modifications. Seeds (25 mg for each genotype) were weighted, and dH_2_O was added. Seeds were shaken using a mixer miller at 30 Hz for 15 min and for other 15 min after turning the samples 180 ° to remove the total mucilage. Seeds were left to settle for 1-2 min. Mucilage-containing supernatant was removed from each sample and transferred to a clean tube. Samples were dried under a stream of N2 gas at 60 °C.

### Alcohol Insoluble solids (AIS) extraction from mature whole seeds

The same seeds from which total mucilage was extracted were used (originally 25 mg), and the extraction was performed according to (Dean et al., 2019). The AIS was destarched in accordance to Zablakis et al. (1995). The resulting AIS was weighted.

### Hydrolysis and Determination of Monosaccharide Composition by HPAEC

Before hydrolysis, a saponification step was performed on the total mucilage and AIS fractions for pectin demethylation. NaOH was added to a final concentration of 250 mM, and samples were incubated for 1 h at RT, before neutralization with 250 mM HCl. Samples were centrifuged for 5 min, supernatants were removed and the pellets were dried. Pellets were then subjected to hydrolysis with trifluoroacetic acid (TFA) as described by Dean et al. (2019). The residual pellet of the TFA hydrolysis of the AIS was in turn hydrolyzed with sulfuric acid (H2SO4) to hydrolyze cellulosic material (Dean et al., 2019).

All the dried hydrolisates were dissolved in ultrapure H_2_O, vortexed, centrifuged and the supernatants transferred to new tubes for high-performance anion-exchange chromatography with pulsed amperometric detection (HPAEC-PAD) analysis. Neutral sugar standards for HPAEC-PAD analysis were prepared and processed as for the total mucilage and AIS. Standard calibration curves were then used to quantify monosaccharide amounts. AIS samples were diluted 1:4. Sample aliquots (10 μl) were analyzed by HPAEC-PAD using a 3 × 150 mm anionic exchange column Carbopac PA20 with guard column (Dionex Thermo Fischer) mounted on a ICS3000 (Dionex Thermo Fischer) HPLC. Before injection of each sample, the column was washed with 200 mM NaOH for 5 min, then equilibrated with 6 mM NaOH for 10 min. Separation was obtained at a flow-rate of 0.4 mL/min applying an isocratic elution with 6 mM NaOH for 20 min then a linear gradient from 6 mM NaOH to 800 mM NaOH for 10 min. Monosaccharides were detected by using a pulsed gold amperometric detector set on waveform A, according to the manufacturer’s instructions. Peaks were identified and quantified by comparison with the standard mixture. Sugars from mucilage were normalized to the mass of seed extracted, and sugars from whole-seed AIS were normalized to the mass of AIS.

### Statistical analysis

Statistical analyses for ANOVA followed by post-hocTukey’s HSD test (95% family-wise confidence level, alpha level 0.05) and for the compact letter display (cld) were performed using the software R 3.6.3 (https://www.r-project.org/) and Rstudio, PBC, Boston, MA (http://www.rstudio.com/). Unpaired Student’s t test was calculated on GraphPad (https://www.graphpad.com/quickcalcs/ttest1.cfm).

### Bioinformatic analyses

Bioinformatic analyses were performed through these following software and database. For genetic and molecular data, the databases The Arabidopsis Information Resource (TAIR) and Uniprot (https://www.uniprot.org/) were used.

For molecules modeling PDB server and the software PyMOL were used (https://www.wwpdb.org/; https://pymol.org/2/). Predicted signal peptide and membrane-spanning regions was studied by the tool SignaIP 4.1 (http://www.cbs.dtu.dk/services/SignalP/). For localization of transcripts was used eFP Browser (Schmid et al., 2005; Winter et al., 2007); https://bar.utoronto.ca/efp/cgi-bin/efpWeb.cgi). Differential expression analyses were performed by Genevestigator (Hruz et al., 2008), though the Arabidopsis RNAseq data and Affymetrix Arabidopsis ATH1 Genome Array provided by the software. For quantification analysis the software ImageJ was used (https://imagej.nih.gov/ij/).

## ACKNOWLEDGEMENTS

The authors gratefully acknowledge the Sapienza Research Infrastructure for the Mass spectrometry analysis at Maldi/TOF–TOF lab Sapienza, funded by the Large Equipment Project 2018 GA118164897F9CB1, and the Microscopy Centre (Department of Life, Health and Environmental Sciences, University of L’Aquila) for support and assistance in confocal and scanning electron microscopy (SEM).

## AUTHOR CONTRIBUTIONS

GDL, FC, SC, MB and BM conceived the project and supervised the research. SC isolated mutant plants and performed the analysis of defense responses, seed coat and mucilage composition. MB performed the enzymatic characterizations and prepared the oxidized oligosaccharides. DP performed the HPLC and MS analysis. MG performed SEM and confocal microscopy analysis. SC, MB, FC and GDL wrote the manuscript draft, whereas FC, BM, MB and GDL edited the final version of the manuscript. All authors have approved the final manuscript.

The authors responsible for distribution of materials integral to the findings presented in this article in accordance with the policy described in the Instructions for Authors (www.plantphysiol.org) are: Manuel Benedetti and Giulia De Lorenzo.

## SUPPLEMENTAL MATERIALS

Figure S1. The predicted model of *At5g44360/ BBE23* with a close-up view of the type IV catalytic site

Figure S2. Oxidizing activity of FHS-CELLOX1 towards MLGs.

Figure S3. TRIB is not a substrate of CELLOX1.

Figure S4. The Mixed-Linkage Glucans (MLGs) that are substrates of CELLOXs have a β-1→4 glycosidic linkage at the reducing end.

Figure S5. Characterization of the T-DNA insertional mutants *cellox1* and *cellox2.*

Figure S6. Rescue of increased susceptibility to B. *cinerea* infection by transgene-complemented *cellox1* knockout.

Figure S7. Differential expression of BBE-like family genes against biotic pertubations.

Figure S8. CELLOX2 is not expressed in seedlings and is not induced upon elicitor treatment.

Figure S9. Localization of the transcripts of *CELLOX2/At5g44360* and *CELLOX1/At4g20860* in the developing seed according to the database eFPbrowser.

Figure S10. Analysis of *CELLOX1* and *CELLOX2 expression* in seeds.

Figure S11. Monosaccharide composition of total mucilage extracts and AIS fractions from de-mucilaged seeds.

Figure S12. Relative monosaccharide composition (mol %) upon H2SO4 hydrolysis of the residue obtained from TFA hydrolysis of the seed AIS.ù

Figure S13. Percentage of germination for each mutant

## REFERENCES

Bain M, van de Meene A, Costa R, Doblin MS (2020) Characterisation of Cellulose Synthase Like F6 (CslF6) mutants shows altered carbon metabolism in β-D-(1,3;1,4)-glucan deficient grain in *Brachypodium distachyon*. Front Plant Sci 11:602850

Barghahn S, Arnal G, Jain N, Petutschnig E, Brumer H, Lipka V (2021) Mixed-linkage β-1,3/1,4-glucan oligosaccharides induce defense responses in *Hordeum vulgare* and *Arabidopsis thaliana*. Front Plant Sci 12:682439

Ben-Tov D, Abraham Y, Stav S, Thompson K, Loraine A, Elbaum R, de Souza A, Pauly M, Kieber JJ, Harpaz-Saad S (2015) COBRA-LIKE2, a member of the glycosylphosphatidylinositol-anchored COBRA-LIKE family, plays a role in cellulose deposition in arabidopsis seed coat mucilage secretory cells. Plant Physiol 167:711–724

Benedetti M, Verrascina I, Pontiggia D, Locci F, Mattei B, De Lorenzo G, Cervone F (2018) Four Arabidopsis berberine bridge enzyme-like proteins are specific oxidases that inactivate the elicitor-active oligogalacturonides. Plant Journal 94:260–273

Burton RA, Collins HM, Kibble NA, Smith JA, Shirley NJ, Jobling SA, Henderson M, Singh RR, Pettolino F, Wilson SM, Bird AR, Topping DL, Bacic A, Fincher GB (2011) Over-expression of specific HvCslF cellulose synthase-like genes in transgenic barley increases the levels of cell wall (1,3;1,4)-β-d-glucans and alters their fine structure. Plant Biotechnol J 9:117–135

Claverie J, Balacey S, Lemaitre-Guillier C, Brule D, Chiltz A, Granet L, Noirot E, Daire X, Darblade B, Heloir MC, Poinssot B (2018) The cell wall-derived xyloglucan is a new DAMP triggering plant immunity in *Vitis vinifera* and *Arabidopsis thaliana*. Front Plant Sci 9:1725

Custers JH, Harrison SJ, Sela-Buurlage MB, van Deventer E, Lageweg W, Howe PW, van der Meijs PJ, Ponstein AS, Simons BH, Melchers LS, Stuiver MH (2004) Isolation and characterisation of a class of carbohydrate oxidases from higher plants, with a role in active defence. Plant J 39:147–160

Daniel B, Konrad B, Toplak M, Lahham M, Messenlehner J, Winkler A, Macheroux P (2017) The family of berberine bridge enzyme-like enzymes: A treasure-trove of oxidative reactions. Arch. Biochem. Biophys. 632:88–103

De Lorenzo G, Cervone F (2022) Plant immunity by damage-associated molecular patterns (DAMPs). Essays Biochem 66:459–469

De Lorenzo G, Ferrari S, Cervone F, Okun E (2018) Extracellular DAMPs in plants and mammals: immunity, tissue damage and repair. Trends Immunol 39:937–950

Dean GH, Sola K, Unda F, Mansfield SD, Haughn GW (2019) Analysis of monosaccharides from Arabidopsis seed mucilage and whole seeds using HPAEC-PAD. Bio Protoc 9:e3464

Di Marzo M, Babolin N, Viana VE, de Oliveira AC, Gugi B, Caporali E, Herrera-Ubaldo H, Martínez-Estrada E, Driouich A, de Folter S, Colombo L, Ezquer I (2022) The Genetic Control of SEEDSTICK and LEUNIG-HOMOLOG in Seed and Fruit Development: New Insights into Cell Wall Control. Plants (Basel) 11

Domon B, Costello CE (1988) A systematic nomenclature for carbohydrate fragmentations in FAB-MS/MS spectra of glycoconjugates. Glycoconjugate Journal 5:397–409

Felle HH, Herrmann A, Hückelhoven R, Kogel KH (2005) Root-to-shoot signalling: apoplastic alkalinization, a general stress response and defence factor in barley (Hordeum vulgare). Protoplasma 227:17–24

Griffiths JS, North HM (2017) Sticking to cellulose: exploiting Arabidopsis seed coat mucilage to understand cellulose biosynthesis and cell wall polysaccharide interactions. New Phytol 214:959–966

Griffiths JS, Tsai AY, Xue H, Voiniciuc C, Sola K, Seifert GJ, Mansfield SD, Haughn GW (2014) SALT-OVERLY SENSITIVE5 mediates Arabidopsis seed coat mucilage adherence and organization through pectins. Plant Physiol 165:991–1004

Huang Z, Boubriak I, Osborne DJ, Dong M, Gutterman Y (2008) Possible role of pectin-containing mucilage and dew in repairing embryo DNA of seeds adapted to desert conditions. Ann Bot 101:277–283

Karimi M, Inze D, Depicker A (2002) GATEWAY vectors for Agrobacterium - mediated plant transformation. Trends Plant Sci 7:193–195

Kim SJ, Zemelis-Durfee S, Jensen JK, Wilkerson CG, Keegstra K, Brandizzi F (2018) In the grass species *Brachypodium distachyon*, the production of mixed-linkage (1,3;1,4)-beta-glucan (MLG) occurs in the Golgi apparatus. Plant J 93:1062–1075

Locci F, Benedetti M, Pontiggia D, Citterico M, Caprari C, Mattei B, Cervone F, De Lorenzo G (2019) An Arabidopsis berberine bridge enzyme-like protein specifically oxidizes cellulose oligomers and plays a role in immunity. Plant Journal 98:540–554

Martín-Dacal M, Fernández-Calvo P, Jiménez-Sandoval P, López G, Garrido-Arandía M, Rebaque D, Del Hierro I, Berlanga DJ, Torres M, Kumar V, Mélida H, Pacios LF, Santiago J, Molina A (2022) Arabidopsis immune responses triggered by cellulose-and mixed-linked glucan-derived oligosaccharides require a group of leucine-rich repeat malectin receptor kinases. Plant J 10.1111/tpj.16088

Mélida H, Bacete L, Ruprecht C, Rebaque D, Del Hierro I, Lopez G, Brunner F, Pfrengle F, Molina A (2020) Arabinoxylan-oligosaccharides act as damage associated molecular patterns in plants regulating disease resistance. Front Plant Sci 11:1210

Mendu V, Griffiths JS, Persson S, Stork J, Downie AB, Voiniciuc C, Haughn GW, DeBolt S (2011) Subfunctionalization of cellulose synthases in seed coat epidermal cells mediates secondary radial wall synthesis and mucilage attachment. Plant Physiol 157:441–453

Messenlehner J, Hetman M, Tripp A, Wallner S, Macheroux P, Gruber K, Daniel B (2021) The catalytic machinery of the FAD-dependent AtBBE-like protein 15 for alcohol oxidation: Y193 and Y479 form a catalytic base, Q438 and R292 an alkoxide binding site. Arch Biochem Biophys 700:108766

Pettolino F, Sasaki I, Turbic A, Wilson SM, Bacic A, Hrmova M, Fincher GB (2009) Hyphal cell walls from the plant pathogen *Rhynchosporium secalis* contain (1,3/1,6)-β-D-glucans, galacto-and rhamnomannans, (1,3;1,4)-β-D-glucans and chitin. FEBS J 276:3698–3709

Pontiggia D, Benedetti M, Costantini S, De Lorenzo G, Cervone F (2020) Dampening the DAMPs: how plants maintain the homeostasis of cell wall molecular patterns and avoid hyper-immunity. Front Plant Sci 11:613259

Pontiggia D, Ciarcianelli J, Salvi G, Cervone F, De Lorenzo G, Mattei B (2015) Sensitive detection and measurement of oligogalacturonides in Arabidopsis. Front Plant Sci 6:258

Puigbo P, Guzman E, Romeu A, Garcia-Vallve S (2007) OPTIMIZER: a web server for optimizing the codon usage of DNA sequences. Nucleic Acids Res 35:W126–131

Raviv B, Aghajanyan L, Granot G, Makover V, Frenkel O, Gutterman Y, Grafi G (2017) The dead seed coat functions as a long-term storage for active hydrolytic enzymes. PLoS One 12:e0181102

Rebaque D, Del Hierro I, López G, Bacete L, Vilaplana F, Dallabernardina P, Pfrengle F, Jordá L, Sánchez-Vallet A, Pérez R, Brunner F, Molina A, Mélida H (2021) Cell wall-derived mixed-linked β-1,3/1,4-glucans trigger immune responses and disease resistance in plants. Plant Journal 106:601–615

Salmeán AA, Duffieux D, Harholt J, Qin F, Michel G, Czjzek M, Willats WGT, Hervé C (2017) Insoluble (1 → 3), (1 → 4)-β-D-glucan is a component of cell walls in brown algae (Phaeophyceae) and is masked by alginates in tissues. Sci Rep 7:2880

Samar D, Kieler JB, Klutts JS (2015) Identification and deletion of Tft1, a predicted glycosyltransferase necessary for cell wall β-1,3;1,4-glucan synthesis in *Aspergillus fumigatus*. PLoS One 10:e0117336

Scafati V, Troilo F, Ponziani S, Giovannoni M, Scortica A, Pontiggia D, Angelucci F, Di Matteo A, Mattei B, Benedetti M (2022) Characterization of two 1,3-β-glucan-modifying enzymes from Penicillium sumatraense reveals new insights into 1,3-β-glucan metabolism of fungal saprotrophs. Biotechnol Biofuels Bioprod 15:138

Schneider CA, Rasband WS, Eliceiri KW (2012) NIH Image to ImageJ: 25 years of image analysis. Nat Methods 9:671–675

Scortica A, Capone M, Narzi D, Frezzini M, Scafati V, Giovannoni M, Angelucci F, Guidoni L, Mattei B, Benedetti M (2021) A molecular dynamics-guided mutagenesis identifies two aspartic acid residues involved in the pH-dependent activity of OG-OXIDASE 1. Plant Physiol Biochem 169:171–182

Scortica A, Giovannoni M, Scafati V, Angelucci F, Cervone F, De Lorenzo G, Benedetti M, Mattei B (2022) Berberine bridge enzyme-like oligosaccharide oxidases act as enzymatic transducers between microbial glycoside hydrolases and plant peroxidases. Mol Plant Microbe Interact 10.1094/MPMI-05-22-0113-TA

Sørensen I, Pettolino FA, Wilson SM, Doblin MS, Johansen B, Bacic A, Willats WG (2008) Mixed-linkage (1→3),(1→4)-β-D-glucan is not unique to the Poales and is an abundant component of *Equisetum arvense* cell walls. Plant J 54:510–521

Sullivan S, Ralet MC, Berger A, Diatloff E, Bischoff V, Gonneau M, Marion-Poll A, North HM (2011) CESA5 is required for the synthesis of cellulose with a role in structuring the adherent mucilage of Arabidopsis seeds. Plant Physiol 156:1725–1739

Toplak M, Wiedemann G, Ulicevic J, Daniel B, Hoernstein SNW, Kothe J, Niederhauser J, Reski R, Winkler A, Macheroux P (2018) The single berberine bridge enzyme homolog of *Physcomitrella patens* is a cellobiose oxidase. FEBS J 285:1923–1943

Tseng YH, Scholz SS, Fliegmann J, Krüger T, Gandhi A, Furch ACU, Kniemeyer O, Brakhage AA, Oelmüller R (2022) CORK1, A LRR-Malectin Receptor Kinase, Is Required for Cellooligomer-Induced Responses in. Cells 11

Voiniciuc C, Schmidt MH, Berger A, Yang B, Ebert B, Scheller HV, North HM, Usadel B, Gunl M (2015) MUCILAGE-RELATED10 produces galactoglucomannan that maintains pectin and cellulose architecture in Arabidopsis seed mucilage. Plant Physiol 169:403–420

Wang M, Xu Z, Ahmed RI, Wang Y, Hu R, Zhou G, Kong Y (2019) Tubby-like Protein 2 regulates homogalacturonan biosynthesis in Arabidopsis seed coat mucilage. Plant Mol Biol 99:421–436

Willats WG, McCartney L, Knox JP (2001) In-situ analysis of pectic polysaccharides in seed mucilage and at the root surface of Arabidopsis thaliana. Planta 213:37–44

Yang C, Liu R, Pang J, Ren B, Zhou H, Wang G, Wang E, Liu J (2021) Poaceae-specific cell wall-derived oligosaccharides activate plant immunity via OsCERK1 during *Magnaporthe oryzae* infection in rice. Nat Commun 12:2178

Zablackis E, Huang J, Muller B, Darvill AG, Albersheim P (1995) Characterization of the cell-wall polysaccharides of Arabidopsis thaliana leaves. Plant Physiol 107:1129–1138

Zang H, Xie S, Zhu B, Yang X, Gu C, Hu B, Gao T, Chen Y, Gao X (2019) Mannan oligosaccharides trigger multiple defence responses in rice and tobacco as a novel danger-associated molecular pattern. Mol Plant Pathol 20:1067–1079

Zhao L, Wang S, Fu YB, Wang H (2019) Arabidopsis Seed Stored mRNAs are Degraded Constantly over Aging Time, as Revealed by New Quantification Methods. Front Plant Sci 10:1764

